# Cellpin enables reference-based imputation and denoising of spatial transcriptomes

**DOI:** 10.64898/2026.06.02.729566

**Authors:** Philipp Putze, Daniele Lucarelli, Deelaka Wellappili, Mojtaba Bahrami, Malte D. Luecken, Fabian J. Theis, Dieter Saur

## Abstract

Spatially resolved transcriptomics enables gene expression profiling within tissue architecture, but targeted panels leave much of the transcriptome unmeasured and spatial artifacts such as RNA diffusion and segmentation errors introduce technical noise. These limitations necessitate computational imputation and denoising, yet existing methods typically incorporate spatial measurements during training, limiting scalability and risking the embedding of technology-specific artifacts into learned representations. To address this, we present cellpin, a variational autoencoder trained exclusively on single-cell RNA sequencing data, using teacher-student latent distillation and noise-simulating augmentations to jointly impute unmeasured genes and denoise spatial profiles without requiring cross-modality alignment. Benchmarked against six methods across multiple paired datasets, cellpin achieves superior held-out gene prediction while scaling efficiently to atlas-size references and multi-sample cohorts. In full-transcriptome Atera data, cellpin reduces residual spatial noise and improves cell-state resolution, providing a scalable and principled foundation for biological discovery from spatial transcriptomics data.

## Introduction

Understanding gene expression framed within the spatial organization of tissues is a fundamental goal of modern biology. Bulk RNA sequencing revealed the molecular composition of tissues but lacked single-cell resolution, obscuring cellular heterogeneity. The advent of single-cell RNA sequencing (scRNA-seq) enabled transcriptome-wide profiling at cellular resolution, transforming our understanding of cellular heterogeneity and gene expression dynamics^1,2^. Spatially resolved transcriptomic technologies, such as Xenium from 10X Genomics and MERSCOPE^3^ from Vizgen, further extended this understanding by capturing the spatial positions of individual cells alongside their transcriptomic profiles. This enabled the mapping of transcriptional programs directly onto tissue architecture^2,4,5^ and contextualized gene expression within the local cellular environment^4,6,7^. However, substantial analytical challenges remain that limit the biological insights that can be drawn from these measurements.

These challenges stem from two intertwined limitations of current platforms. First, commercially available panels typically measure only a few hundred genes, leaving most of the transcriptome unobserved. While subsequent generations expand the number of measured genes, they usually yield more sparse readouts^8^. Second, expression measurements are inherently noisy due to technical artifacts, including sparse transcript abundance, imperfect cell segmentation^9,10,11^, and diffusion of free-floating RNA, which can incorrectly assign transcripts to neighbouring cells^8^. Emerging platforms, like Atera (10X Genomics), now offer whole-transcriptome spatial profiling, representing the current state-of-the-art in spatial resolution and transcriptomic coverage; however, the effect of spatial noise and artifacts on the resolution of this new technology remains unclear. These noise sources can blur the expression signatures of molecularly similar populations, limiting the resolution of downstream analyses, and identification of biologically relevant cellular lineages and states distinguished by subtle transcriptomic differences.

Imputation, the prediction of unmeasured gene expression by using orthogonal datasets, offers a principled approach to recovering transcriptome-wide coverage, leading to the development of a diverse set of computational approaches^12,13^. Methods such as Tangram^14^ and gimVI^15^ align spatial and single-cell data in a shared latent space to transfer expression profiles. SpaGE^16^ leverages a shared latent representation using domain adaptation to project between modalities. Despite their utility, most existing approaches are constrained by a common conceptual assumption: that spatial and single-cell measurements should be aligned during model training. In practice, enforcing this alignment fails when a cell state is present in one modality but missing from the other, as the model will erroneously force that unique state to match an incorrect cell type. Directly incorporating spatial data into training may also propagate modality-specific technical artifacts, like transcript diffusion and segmentation errors, into the learned representation^17^, and attempting to learn a corrected representation from the spatial data itself creates a circular dependency, where the model inherently encodes and propagates the very technical biases they are designed to remove. This may limit the resolution of rare or transcriptionally similar populations. At the same time, joint training over growing numbers of spatial samples poses substantial computational scalability challenges and complicates atlas-scale, multi-sample analysis.

We developed cellpin to address these limitations: a lightweight probabilistic framework that learns a biologically meaningful latent representation from scRNA-seq references alone. Cellpin uses this representation to denoise spatial expression profiles, recover latent structure, and enable transcriptome-wide imputation from sparse spatial measurements at scale. It requires no paired spatial measurements and transfers directly across datasets without retraining, explicitly accommodating the technical characteristics of spatial measurement without imposing a one-to-one mapping between single-cell and spatial data.

Applied to the nascent Atera platform, cellpin improved on the state-of-the-art transcriptomic coverage by enhancing downstream analyses through substantially denoised expression profiles. Together, these features establish cellpin as a unified and scalable approach for spatial transcriptomics analysis. We envision it as a broadly applicable tool for unlocking biological insights from both current panel-based and emerging full-transcriptome spatial datasets.

## Results

### Cellpin imputes unmeasured genes combining generative modelling and latent-space distillation

The cellpin framework is implemented as a two-stage variational autoencoder trained exclusively on scRNA-seq data, such that no spatial coordinates or matched spatial measurements are required during model fitting, hence avoiding spatial measurement noise and biases. (**Fig. 1**; see **Methods**). In the first stage, a teacher model learns to reconstruct full-gene scRNA-seq profiles, establishing a structured latent representation of cellular transcriptional states. In the second stage, a student encoder is trained to infer the same latent structure from panel genes alone while preserving accurate full-transcriptome reconstruction through a shared decoder. This second stage is guided jointly by reconstruction losses, which preserve gene-level fidelity, and latent-matching losses, which encourage the panel-based representation to remain consistent with the full-transcriptome latent space (**Supplementary Fig. 1 a-b**).

**Figure 1:**
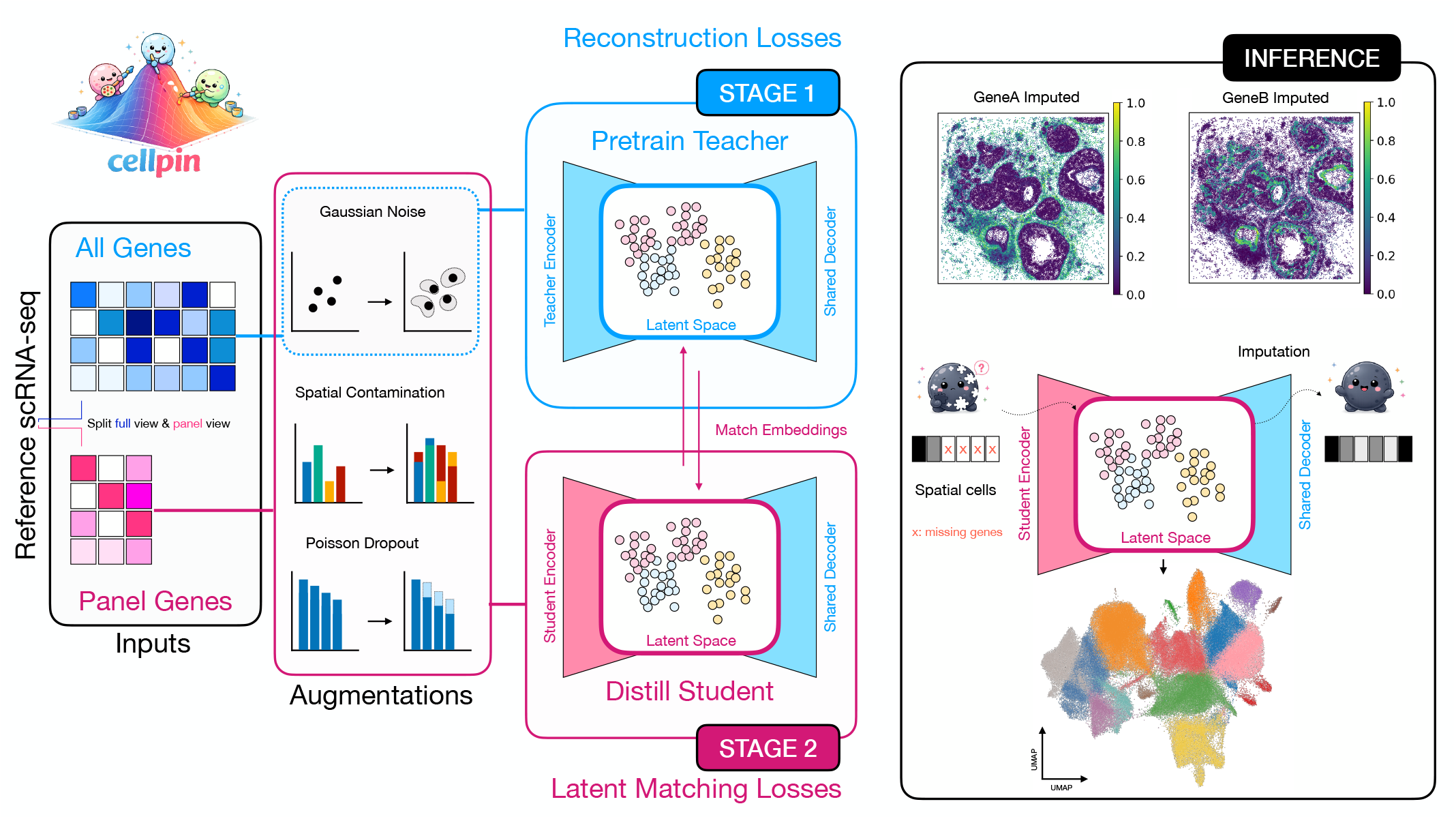
Cellpin aligns full-gene and panel-gene representations for robust spatial imputation. Cellpin is trained on a two-stage process. In the first stage (blue), the reference single-cell dataset is used to train a Teacher Variational Auto Encoder (VAE)^19^ with reconstruction losses. At this stage, the count data is only augmented with Gaussian Noise. After the first stage is done, the model enters a second stage of training (pink), where the cells are split into two views: one with all the genes to reconstruct, and a second one with only the genes present in the spatial dataset to impute. During this phase, the student encoder is forced to learn a representation of the reduced view as close as possible to the full view, using contrastive losses (see **Methods**). The views are augmented with Poisson dropout and transcript contamination to simulate missegmentation and other spatial artifacts. After the model is trained, it can be used to impute a spatial dataset and extract meaningful embeddings from it (black).

Because cellpin is trained only on dissociated scRNA-seq reference cells, the model learns to reconstruct comparatively clean cellular expression states rather than directly fitting sparse and spatially corrupted measurements. Inspired by the success of previous data augmentation techniques^18^, robustness to domain shift between scRNA-seq and spatial data is explicitly introduced during training via perturbations that mimic reduced capture efficiency, local contamination, and measurement noise (**Fig. 1**; see **Methods**). Once trained on an appropriate reference from the same tissue, biological context, or disease setting, the model can then be applied to new spatial samples measured with the same panel without retraining on each sample individually.

### Cellpin outperforms existing imputation methods across diverse spatial transcriptomics datasets

To evaluate cellpin’s imputation performance, we benchmarked it against several existing spatial transcriptomics imputation methods, including Tangram^14^, gimVI^15^, SpaGE^16^, scENVI^20^, stDiff^21^, and scConcept^22^ in both zero-shot and fine-tuned (scConcept+) settings. While several foundational models have been recently developed, we only benchmarked against scConcept, which has been shown to be the best performer in the spatial imputation task^22^. We performed a five-fold held-out gene prediction using paired single-cell^23–27^ and spatially resolved transcriptomics ^24,25,28^ datasets (**Supplementary Table 1**) spanning diverse cell types, tissue origins, and transcriptional states (**Fig. 2a**). In each fold, a randomly selected set of 50 genes was withheld and used as ground-truth targets for imputation evaluation, while the remaining genes were treated as the observed spatial panel. Gene panels were sampled such that no gene appeared in more than one fold per dataset, ensuring complete independence between benchmarking iterations, with 1250 total genes analyzed for the benchmarking. To comprehensively evaluate imputation performance, we assessed several complementary metrics: Pearson correlation and Lin’s concordance correlation coefficient (CCC), which capture linear agreement and combined precision and accuracy between predicted and measured expression, respectively, with higher values indicating better performance; and root mean square error (RMSE) and Jensen-Shannon divergence (JS), which quantify magnitude of prediction error and divergence between predicted and true expression distributions, respectively, where lower values reflect superior reconstruction (**Fig. 2a**).

**Figure 2:**
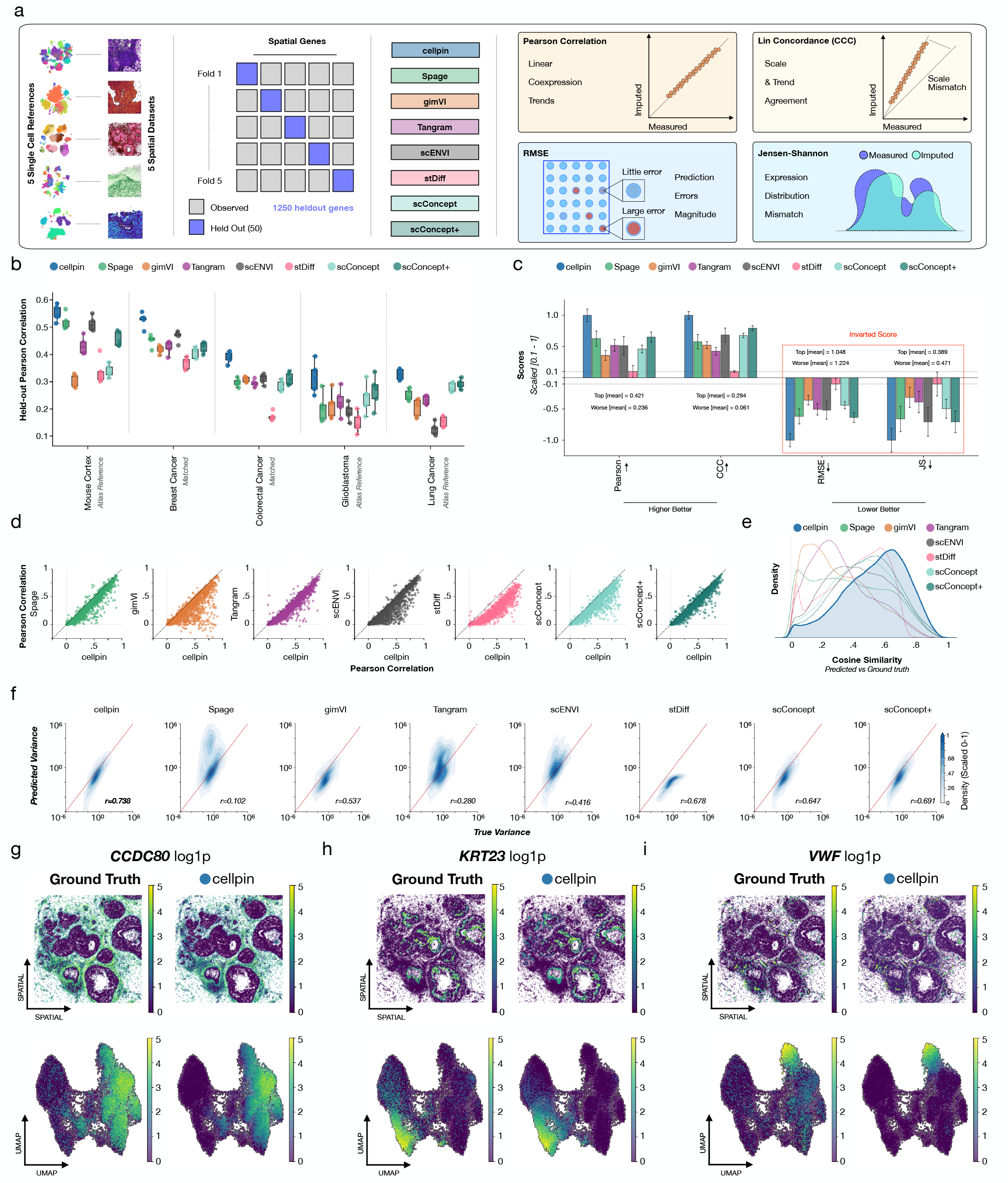
Cellpin outperforms state-of-the-art spatial expression imputation methods across multiple datasets and metrics. **a)** Overview of the benchmarking strategy. Five single-cell reference datasets and five spatial transcriptomics datasets (**Supplementary Table 1**) were used to evaluate cellpin against six state-of-the-art imputation methods. For each spatial dataset, genes were partitioned in a 5-fold cross-validation scheme, with 50 held-out genes per fold (250 held-out genes per dataset total). Methods were tasked with imputing held-out spatial genes from the remaining observed genes, and performance was evaluated across four complementary metrics: Pearson correlation, Lin’s concordance correlation coefficient (CCC), root mean square error (RMSE) and Jensen-Shannon divergence (JS). **b)** Mean held-out Pearson correlation across five spatial datasets, evaluated using matched same-study data and external atlas references. Each dataset was assessed by 5-fold cross-validation with 50 unique held-out genes per fold (250 genes per dataset total). **c)** Summary performance across all benchmarked datasets. Pearson correlation and CCC were min–max scaled (higher=better), whereas RMSE and JS divergence were inverted prior to scaling (lower=better). Bars represent the mean scaled score across all datasets and cross-validation folds; error bars denote the standard error of the mean. Unscaled mean for the top and worst performing method depicted for each metric. **d)** Pairwise comparison of per-gene Pearson correlations between cellpin and competing methods across 1,250 held-out genes. Each point represents one gene, plotted by Pearson correlation between imputed and ground-truth expression for cellpin (x-axis) and the corresponding comparison method (y-axis). Points below the diagonal indicate genes for which cellpin achieved higher predictive performance. **e)** Density distributions of cosine similarity between ground-truth and imputed whole-cell expression vectors across 250 held-out genes per dataset. **f)** True versus predicted expression variance for all held-out genes, with Pearson correlation (*r*) between true and predicted variance annotated per method. **g–i)** Representative expression maps for three held-out genes from a breast cancer Xenium dataset^24^: *CCDC80* (r = 0.86) (g), *KRT23* (r = 0.82) (h), and *VWF* (r = 0.84) (i). For each gene, ground-truth measurements are shown alongside cellpin imputation in tissue spatial coordinates (top) and in UMAP coordinates derived from ground-truth PCA (bottom).

Cellpin consistently achieved the highest held-out gene Pearson correlation across all evaluated methods and independent datasets (**Fig. 2b**). When performance metrics were averaged across datasets, cellpin showed a uniform advantage over all competing approaches, achieving the highest Pearson correlation and CCC, while also yielding the lowest RMSE and Jensen–Shannon divergence (**Fig. 2c, Supplementary Fig. 2a**). These results indicate that cellpin does not merely improve a single accuracy metric but provides a broadly superior reconstruction of spatial transcriptomes across complementary measures of correlation, concordance, error and distributional similarity.

Gene-level benchmarking further confirmed the robustness of this improvement. Cellpin-imputed genes showed higher Pearson correlations not only for global expression trends, but across nearly all individuals held-out genes when compared with state-of-the-art models (**Fig. 2d**). This gene-wise consistency is important because it demonstrates that cellpin’s advantage is not driven by a small subset of highly expressed or easily predictable genes, but reflects a generalizable capacity to reconstruct diverse transcriptional programs from sparse spatial panels.

To assess whether cellpin preserves the high-dimensional structure of gene expression, we next evaluated cosine similarity between predicted and measured expression vectors. Unlike scalar error metrics, cosine similarity captures the alignment of transcriptional profiles in high-dimensional space and therefore reflects the preservation of relative co-expression patterns rather than absolute expression magnitude alone. Cellpin achieved consistently higher cosine similarity across datasets (**Fig. 2e**), indicating that its imputations retain biologically meaningful transcriptional geometry. Importantly, cellpin also showed the strongest agreement between true and imputed gene-level variability (r = 0.738) as well as mean (r = 0.817) among all evaluated methods (**Fig. 2f, Supplementary Fig. 3a**), demonstrating that the model captures fine-grained biological heterogeneity without excessive smoothing.

We next evaluated whether this accuracy is achieved at the cost of computational complexity. Runtime and memory profiling across increasing dataset sizes showed that cellpin was consistently faster while being memory-efficient during both training and inference (**Supplementary Table 2, Supplementary Fig. 3b**). Moreover, training remained robust at scale; held-out gene Pearson correlation declined only marginally as the number of reference genes increased (**Supplementary Table 3**). Thus, cellpin combines improved predictive accuracy with computational scalability, supporting its use for large spatial transcriptomic studies and repeated deployment across datasets without dataset-specific retraining.

Qualitative inspection, using examples from a breast cancer Xenium dataset^24^, confirmed the quantitative benchmarking results. Cellpin reconstructed spatial expression patterns directly from its learned representation for held-out genes spanning a full range of expression levels, from broadly distributed (*CCDC80*) to moderately expressed (*KRT23*) to sparsely detected (*VWF*), faithfully recovering the ground-truth spatial signal in each case (**Fig. 2g-i**).

Lastly, the choice and quality of the single-cell reference dataset used to train cellpin is of paramount importance. When benchmarking the imputation of the same spatial dataset using references generated by different single-cell technologies (10x Chromium and 10x Flex from matched samples), we observed that imputation performance varied drastically based on the reference type (**Supplementary Fig. 3c**). This stark contrast highlights that the biological fidelity and sequencing depth of the data directly constrain the model’s predictive performance.

### Cellpin imputes biologically relevant transcriptional information in large spatial cohorts

To evaluate the ability of cellpin to train once on an atlas-scale reference and deploy across an entire spatial cohort without sample-level matching, we applied a single model to 45 Xenium samples spanning both healthy donors and pulmonary fibrosis (PF) patients, comprising approximately 1.6 million cells^29^, after training it on a subset of the Human Lung Cell Atlas (HLCA)^30^ likewise spanning healthy and PF donors (**Fig. 3a, Supplementary Table 1**). This transfer across distinct cohorts and unmatched donors represents a stringent test case, requiring cellpin to map complex transcriptional architecture without dataset-specific retraining or sample pairing.

**Figure 3:**
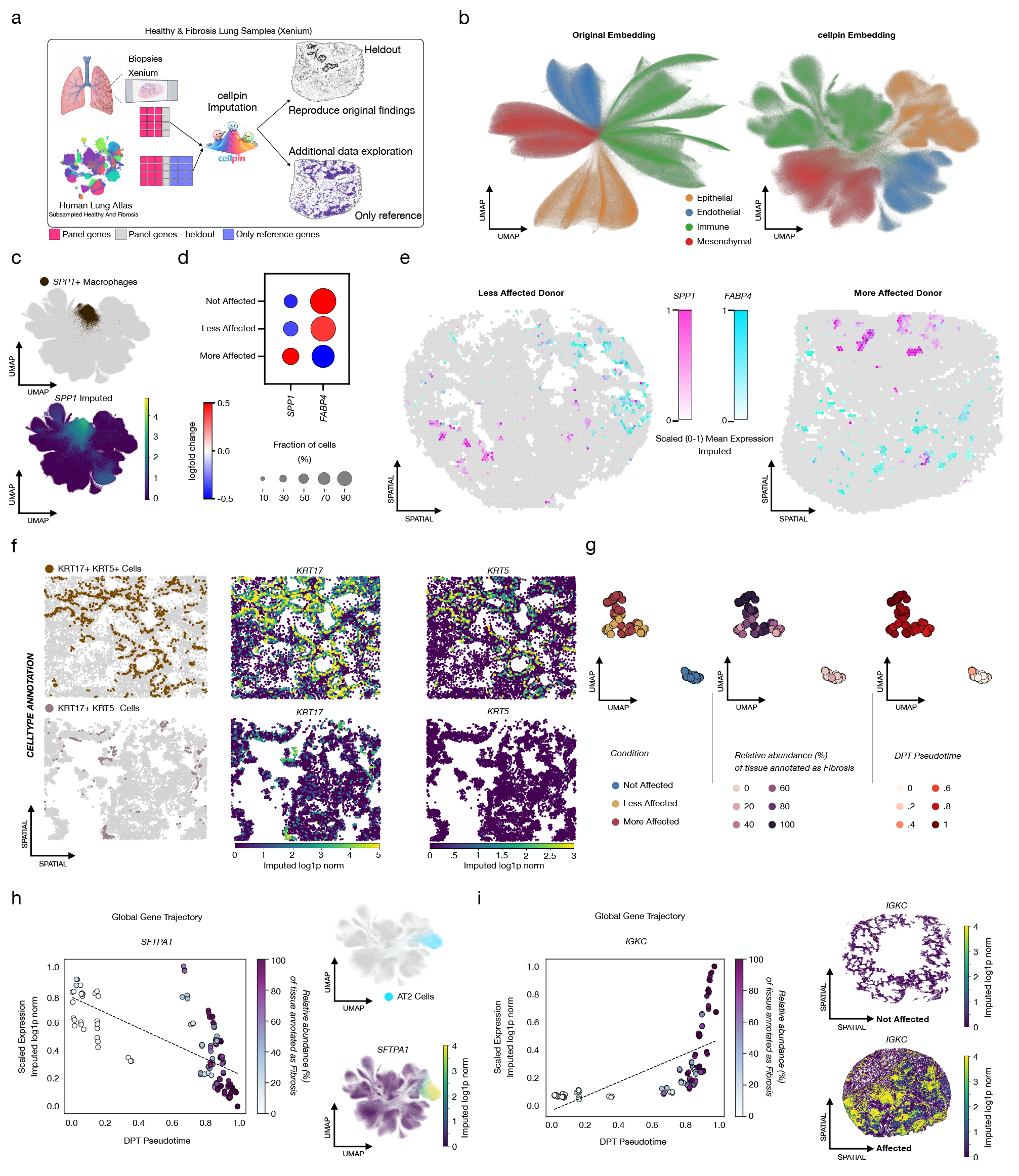
Cellpin recapitulates hallmark pulmonary fibrosis niches and reveals disease trajectory-associated genes from exclusively imputed spatial expression across 45 Xenium samples. **a)** Schematic of the validation strategy. Cellpin was trained on a subsampled subset of the Human Lung Cell Atlas^30^ comprising healthy and idiopathic pulmonary fibrosis donors. Four genes central to the findings of the original study (*KRT5, KRT17, SPP1, FABP4*) were deliberately withheld from the Xenium panel during training and subsequently recovered by imputation. **b)** Uniform manifold approximation projection (UMAP) of the 45 Xenium samples derived from the original study (left) alongside a cellpin embedding-derived UMAP of the same spatial cells (right), colored by major cellular lineage. **c)** Cellpin UMAP of the spatial dataset highlighting *SPP1*+ macrophages as annotated in the original study (top) and corresponding cellpin-imputed *SPP1* expression (bottom). **d)** Differential expression analysis on the cellpin-imputed transcriptome in macrophages, stratified by fibrosis severity annotation from the original study (Not Affected, Less Affected, More Affected). **e)** Representative spatial plots from a less affected and a more affected pulmonary fibrosis donor showing mutually exclusive spatial localization of imputed *SPP1* and *FABP4* expression. Values shown are scaled mean expressions across aggregated spatial bins. **f)** Spatial maps of KRT17+ KRT5+ (basal cells) and KRT17+ KRT5- (aberrant basaloid cells) annotations from the original study (left) alongside cellpin-imputed expression of *KRT17* and *KRT5* (middle and right panel). Both genes were withheld during training. **g)** Patient-level embeddings derived by mean aggregation of per-cell cellpin embeddings using patpy^32^, with three random-cell pseudo-replicates per sample. Embeddings are colored by disease severity (left), percentage of tissue annotated as fibrotic by the study pathologist (middle), and diffusion pseudotime (DPT, right). **h)** Scatter plot of scaled mean pseudoreplicate-level imputed *SFTPA1* expression along the DPT trajectory, colored by pathology annotation, showing progressive downregulation with disease severity. **i)** Scatter plot of scaled mean pseudoreplicate-level imputed *IGKC* expression along the DPT trajectory, colored by pathology annotation, alongside representative spatial maps from an unaffected normal and a more affected fibrotic sample, demonstrating progressive upregulation of *IGKC* with disease severity.

Despite this substantial mismatch, cellpin generated a well-structured latent embedding that resolved major cell populations more clearly than standard dimensionality reduction of raw counts (**Fig. 3b**). This demonstrates a key innovation of cellpin: rather than relying on paired spatial training data, cellpin can use an unpaired single-cell atlas as a reusable reference to denoise and enrich large-scale spatial transcriptomic data across cohorts, technologies and disease contexts.

To directly test biological fidelity, we withheld four genes central to the original study’s conclusions, *SPP1, FABP4, KRT5*, and *KRT17*^*29*^. Cellpin-imputed profiles accurately recapitulated the reported spatial segregation of SPP1+ and FABP4+ alveolar macrophage populations into distinct tissue compartments, with *SPP1* signal correctly concentrated in regions of active tissue destruction (**Fig. 3c-e, Supplementary Figure 4a-b**). Importantly, cellpin also recovered the rare KRT17+/KRT5-aberrant basaloid cell population identified in the original study^29^ which plays an important role in proinflammatory signalling and disease progression^31^. Although both defining marker genes were withheld from the observed panel, cellpin correctly imputed their expression in epithelial cells that had been annotated as this population in the original study based on those same marker genes (**Fig. 3f**).

We next asked whether cellpin embeddings could recover disease-associated progression at the donor level. Pseudobulk cellpin embeddings, constructed by averaging cell embeddings for each donor using patpy^32^, were ordered along a pseudotime axis rooted at a healthy reference donor. This trajectory broadly recapitulated pathologist-assessed fibrosis severity (**Fig. 3g**), indicating that cellpin preserves clinically relevant disease structure across individuals. Genes correlating with this trajectory included two genes absent from the measured spatial panel: *SFTPA1*, a canonical AT2 marker, and *IGKC*, a plasma cell marker. Cellpin-imputed *SFTPA1* expression negatively correlated with disease progression, while *IGKC* expressing plasma cells showed positive correlation, consistent with previous reports^33–35^ (**Fig. 3h-i**), showing the ability of cellpin to reconstruct disease-associated transcriptional trajectories beyond the limits of the measured spatial panel.

### Cellpin denoises spatial profiles, enabling accurate atlas-based label transfer

Cellpin enables not only accurate gene imputation, but also robust downstream biological interpretation through accurate label transfer from a reference atlas. Following training on scRNA-seq data exclusively, cellpin extracts latent embeddings independently for both the single-cell reference and the spatial query dataset. Since the reference atlas provides curated cell type annotations, a lightweight classifier can be trained on single-cell cellpin embeddings to learn the mapping from the latent representation to cell identity. This trained classifier can then be applied directly to the spatial cellpin embeddings, enabling cell type label transfer onto the spatial dataset without any additional model training, spatial fine tuning, or exposure to spatial measurements and corresponding artifacts (**Fig. 4a**).

**Figure 4:**
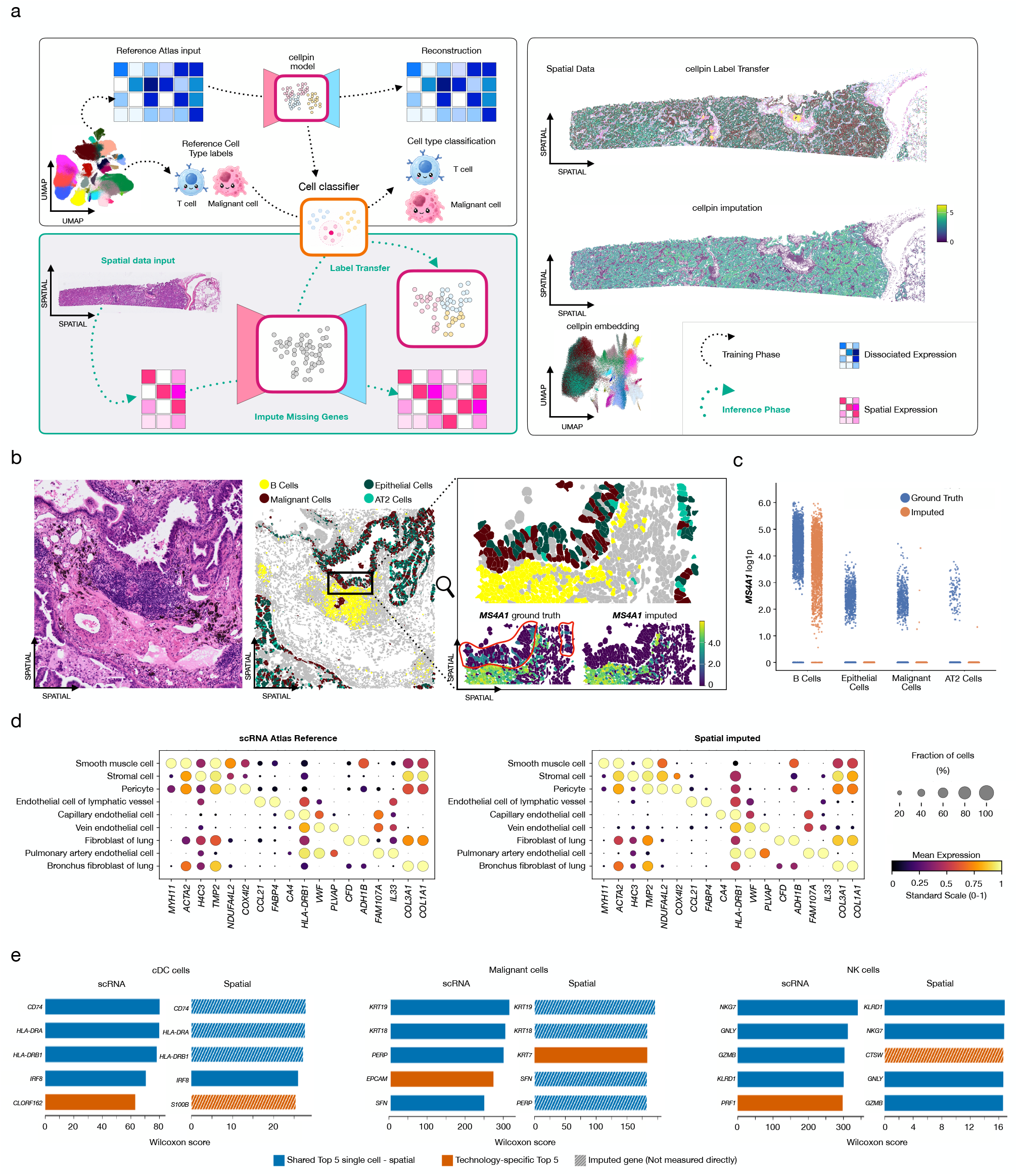
Cellpin enables atlas-guided spatial cell type annotation, denoising, gene expression imputation, and embedding in a single forward pass. **a)** Schematic overview of the cellpin framework. In the first stage, cellpin is trained on a human lung cancer scRNA-seq reference atlas to learn a low-dimensional cell embedding and reconstruct retained gene expression. In a second stage, a cell type classifier is trained on the resulting cellpin embeddings using reference atlas annotations. At inference, a single forward pass maps Xenium spatial transcriptomics data through the panel encoder, simultaneously yielding cell type annotations, expression imputation, and a low-dimensional cell embedding. **b)** Hematoxylin and eosin (H&E)-stained lung tissue section (left) with corresponding cellpin cell type annotations highlighting B cells, malignant cells, epithelial cells and AT2 cells (middle). Zoomed insets show measured and cellpin-imputed expression of the B cell marker *MS4A1* (right). **c)** Per-cell *MS4A1* expression across B cells, epithelial cells, malignant cells, and AT2 cells, comparing measured (blue) and cellpin-imputed (orange) values. **d)** Dot plots showing mean expression of stromal cell type marker genes in the scRNA-seq reference atlas (left) and cellpin-imputed Xenium data (right). Marker genes were identified by differential expression analysis in the reference atlas. **e)** Top differentially expressed genes per cell type (cDC cells, malignant cells, NK cells) in the scRNA-seq reference and cellpin-imputed spatial data, ranked by Wilcoxon score computed independently per modality. Blue bars indicate genes ranking in the top 5 in both modalities; orange bars indicate markers present in the top 5 of one modality only; hatched bars denote genes not measured directly in the Xenium panel and recovered exclusively through cellpin imputation.

To evaluate cellpin’s denoising and label-transfer capabilities, we performed imputation and cell type label transfer (**Supplementary Figure 5a**) on a spatial Xenium dataset of a lung tumor (FFPE sections of Human Lung Cancer profiled with Xenium Multimodal Cell Segmentation) after training on an unmatched lung cancer single-cell atlas as reference^27^ (**Fig. 4a, Supplementary Table 1**). Initial inspection of the measured spatial counts revealed apparent contamination or signal leakage in the ground-truth data. For example, expression of the B cell marker *MS4A1* was detected in a region composed primarily of epithelial, malignant and AT2 cells located adjacent to a B-cell-rich area (**Fig. 4b**). Following cellpin imputation, this region no longer displayed putative ectopic B cell marker expression, but instead showed expression profiles consistent with their expected epithelial, malignant or AT2 identities (**Fig. 4b-c**), demonstrating successful denoising by cellpin due to the training on only single-cell reference data.

Cellpin-imputed marker gene profiles from spatial datasets closely recapitulate those observed in the scRNA-seq reference, including well-separated stromal cell populations (**Fig. 4d**), and additional cellular compartments (**Supplementary Figure 5b-c**). The comparison of top-ranked marker genes across representative cell types, including cDCs, malignant cells, and NK cells, demonstrated strong concordance between the scRNA-seq reference and cellpin-imputed spatial profiles (**Fig. 4e**). In the original spatial panel, *IRF8* was the only top-ranked marker for cDC identity. After cellpin imputation, canonical antigen-presentation markers including *CD74, HLA-DRA*, and *HLA-DRB1* were recovered, substantially enriching the gene signature available for cDC cell type annotation (**Fig. 4e**).

Cellpin similarly improved marker support for malignant and NK cell populations. For malignant cells, imputed expression of *KRT19, KRT18, SFN*, and *PERP* emerged as the defining top-ranked marker genes, enabling robust identification of this population that lacked clear marker support in the measured spatial panel. NK cell identity was similarly well-preserved, with *KLRD1, NKG7, GNLY*, and *GZMB* consistently top-ranked after imputation (**Fig. 4e**). Together, these results support the fidelity of cellpin’s imputation and label-transfer across diverse lineages, recovering biologically coherent and annotation-informative gene signatures that were absent or incomplete in the original spatial measurement.

### Cellpin denoises full-transcriptome Atera spatial data, improving data quality and downstream analyses

The 10x Genomics Atera single-cell spatial transcriptomics platform enables state-of-the-art, high-resolution, whole-transcriptome spatial profiling and substantially expands transcriptome coverage beyond targeted-panel platforms like Xenium. Because Atera is an emerging technology and public performance benchmarks remain limited, we used this modality to test whether cellpin can further improve data quality and downstream biological interpretation in an already information-rich spatial whole transcriptomic setting. We applied cellpin to the preview Atera spatial breast cancer gene expression dataset (10x Atera in Situ Gene Expression, FFPE Human Breast Cancer) after training on an independent single-cell breast cancer atlas^36^ (**Fig. 5a, Supplementary Table 1**).

**Figure 5:**
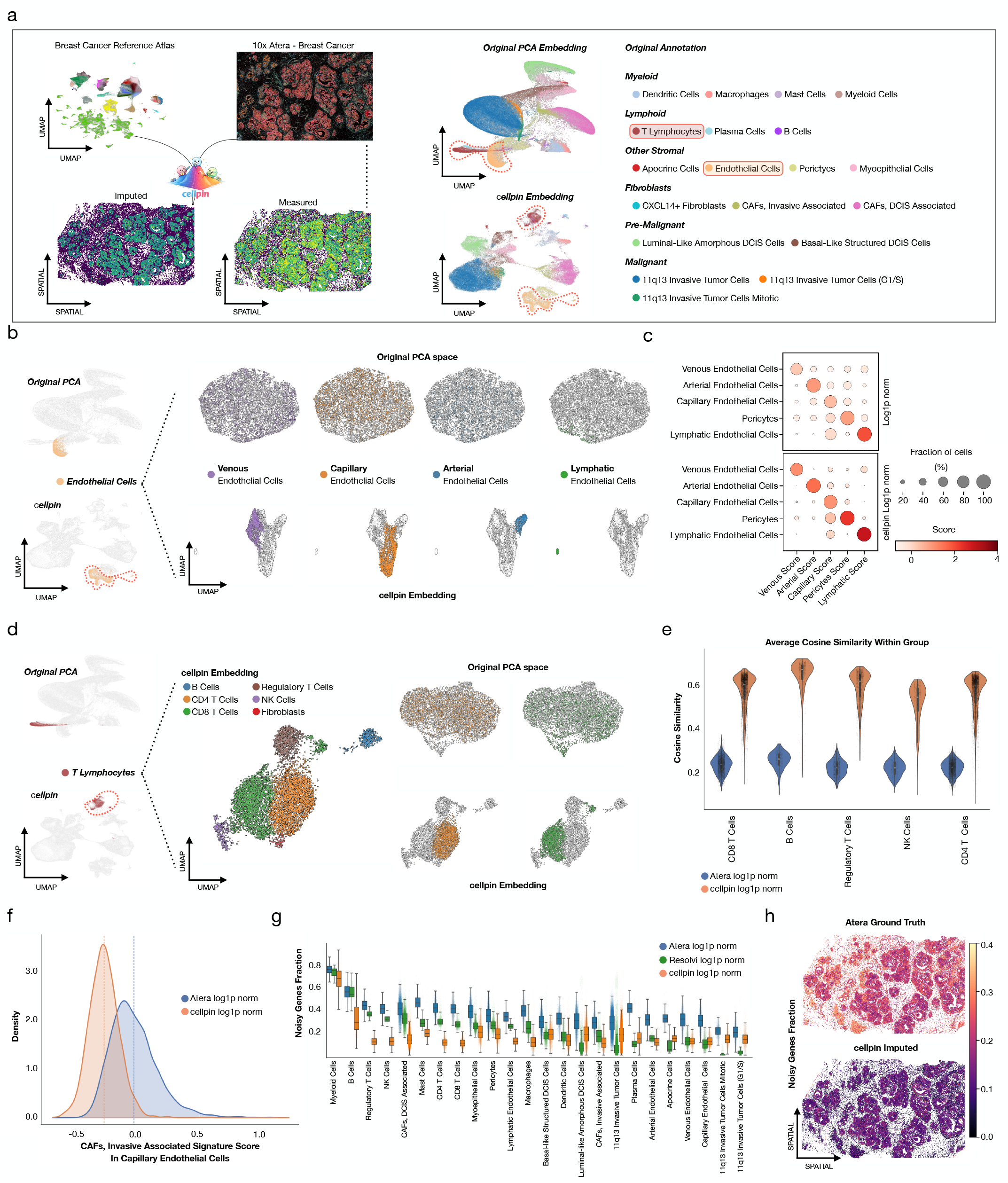
Cellpin improves sensitivity and enables refined downstream analyses in 10x Genomics Atera whole transcriptome single-cell spatial transcriptomics data. **a)** Cellpin was applied to a 10x Genomics Atera (whole-transcriptome single-cell spatial transcriptomics technology) preview dataset of breast cancer tissue, using an independent breast cancer single-cell reference atlas for training. Uniform Manifold Approximation Projection (UMAP) derived from the original PCA on the measured Atera gene expression profiles (top) and from the cellpin latent embedding (bottom) are shown, colored by annotated cell type. **b)** Focused analysis of endothelial cells revealed unresolved clusters in the original PCA embedding of the measured Atera profiles (top). The cellpin embedding separates endothelial cells into four transcriptionally distinct states: venous, capillary, arterial, and lymphatic endothelial cells (bottom). **c)** Dot plots of endothelial subtype signature scores across inferred endothelial cell states, computed on markers derived from rank-based differential expression. Scores are shown for the measured Atera gene space (top) and the cellpin-imputed gene space (bottom). **d)** Focused analysis of T and NK lymphocytes. PCA of the measured Atera profiles shows limited separation of CD4 T cells, CD8 T cells, regulatory T cells and NK cells, with two additional clusters initially annotated as T lymphocytes (top). Cellpin embeddings resolve these populations more clearly and reveal that the two ambiguous T-cell-associated clusters correspond to B cells and fibroblasts based on imputed lineage-marker expression (**Supplementary Figure 6d-e**). **e)** The average cosine similarity for each lymphocyte subtype, computed between each cell and all other cells assigned to the same cluster. Higher values indicate improved transcriptional coherence within inferred cell states. **f)** Distribution of invasive-associated CAF signature scores in capillary endothelial cells, computed from measured Atera and cellpin-imputed profiles. **g)** The fraction of detected genes that are improbable for a cell’s annotated cell type, used as a per-cell estimate of transcriptional noise or ectopic lineage-marker contamination (see methods). Comparing cellpin imputed gene expression (orange) with resolVI (green) and Atera ground truth (blue). **h)** Spatial maps of the noise gene fraction computed from Atera ground-truth measured (top) and cellpin-imputed (bottom) profiles.

Cellpin generated a more structured embedding of the Atera spatial transcriptomic data compared to the original PCA embedding of the measured expression profiles, with improved separation of major cell populations (**Fig. 5a**). To illustrate the downstream impact of this improvement, we focused on two representative lineages, endothelial cells and T lymphocytes, as proof-of-concept examples. A comprehensive re-annotation of the full dataset was beyond the scope of this study. In endothelial cells, PCA of ground-truth-measured spatial counts failed to resolve meaningful substructure, yielding largely overlapping distributions without clear subtype separation. In contrast, cellpin imputation and extraction of cell embeddings enabled the identification and annotation of distinct endothelial lineages, including venous, capillary, arterial, and lymphatic cells (**Fig. 5b**). Cellpin-reconstructed marker genes exhibited substantially reduced noise compared with Atera raw counts, with cleaner separation between marker-positive and marker-negative populations across inferred endothelial cell states (**Fig. 5c, Supplementary Figure 6a-b**). Interestingly, if PCA is computed over the imputed gene space rather than the raw measurements, the same cell states can be recovered (**Supplementary Figure 6c**).

A similar improvement was observed in T lymphocytes. Cellpin recovered transcriptionally distinct T cell subpopulations that were not apparent in the measured spatial profiles (**Fig. 5d**). Moreover, using cellpin embeddings we were able to correct an apparent annotation artifact in the original dataset, in which B cells and fibroblasts were partially mixed with T lymphocytes (**Fig. 5d, Supplementary Figure 6d-e**). To quantify the resulting improvement in transcriptional coherence, we computed within cell type cosine similarity across lymphocyte subsets. Across all major subtypes, cellpin imputed profiles exhibited higher intra-class similarity than the measured ground-truth spatial data, indicating reduced technical noise and improved consistency in cell type–specific expression programs (**Fig. 5e**). These examples demonstrate that cellpin denoising enhances the biological signal by suppressing variability introduced by spatial measurement artifacts while preserving lineage-specific transcriptional structure.

To directly quantify noise reduction, we next computed a cancer-associated fibroblast (CAF) invasive associated signature score using both measured ground-truth and cellpin-reconstructed profiles. In the raw spatial profiles, capillary endothelial cells exhibited relatively elevated CAF scores, consistent with their spatial proximity to CAFs and as observed following spatial neighborhood analysis (**Supplementary Figure 6f**). After cellpin imputation, the CAF signal in capillary endothelial cells was substantially reduced, indicating improved separation of neighboring but transcriptionally distinct cell types (**Fig. 5f**).

To generalize this principle, and further quantify transcriptional contamination, we defined a noisy gene fraction per cell, capturing the proportion of genes with high likelihood of ectopic expression in unrelated cell types, such as lineage markers detected outside their expected compartment (e.g., B cell genes in T cells, see **Methods**). Across the tissue section, cellpin markedly reduced the noisy gene fraction relative to ground-truth spatial Atera as well as ResolVI-corrected^37^ imputed measurements, indicating improved specificity of gene expression signals (**Fig. 5g**). Spatial mapping of this metric further revealed that noise reduction was consistent across tissue regions rather than restricted to specific compartments (**Fig. 5h**), altogether demonstrating the ability of cellpin to complement next-generation full-transcriptome spatial technologies by enhancing the biological resolution and interpretability of complex information-rich spatial datasets.

## Discussion

Cellpin is a probabilistic approach that addresses two fundamental limitations of spatial transcriptomics: the panel bottleneck, which restricts commercial platforms to a fraction of the transcriptome, and technical noise arising from common spatial artifacts, like sparse transcript abundance and RNA diffusion. By training exclusively on scRNA-seq data and incorporating physically inspired spatial augmentations, cellpin achieves robust cross-modal transfer without requiring matched spatial-single-cell pairs. Across diverse benchmarks, cellpin outperformed existing imputation methods in held-out gene prediction, recovering both widespread and rare spatial expression patterns with greater fidelity. These gains translated directly into downstream analytical improvements: identifying biological context from large cohorts of lung fibrosis; more accurate cell type label transfer and denoising of contaminated expression profiles in lung cancer tissue; and enhanced resolution of transcriptional subpopulations in full-transcriptome Atera breast cancer data. This demonstrated that imputation and denoising can be unified within a single lightweight inference framework.

Cellpin distinguished itself from existing imputation methods like Tangram, gimVI and SpaGE by learning a generative model of transcriptional states from a scRNA-seq reference alone, with simulated spatial artifacts during training, rather than explicitly mapping between paired spatial and single-cell datasets. This sidesteps the need for dataset-specific alignment entirely and naturally scales to large atlas references that provide a demonstrably robust and generalizable single-cell reference but would challenge the current alignment-based approaches. At inference, a Monte Carlo dropout strategy further improves robustness to stochastic gene missingness in sparse spatial panels, at no additional training cost. Cellpin is also fully compatible with the scverse^38^ ecosystem, accepting inputs and returning outputs that slot directly into anndata^39^, and spatial data^40^, making it easy to integrate with the broader suite of scverse tools, requiring no additional data wrangling to fit into existing single-cell and spatial analysis workflows.

The denoising behaviour of cellpin is perhaps its most practically significant property. Prior methods that incorporate spatial measurements during training risk directly absorbing artifacts into their learned representations. By contrast, cellpin acts as a denoising and representation-learning layer, with its single-cell generative prior serving as a clean reference that suppresses ectopic marker expression without ever having seen spatial data corrupted by artifacts such as missegmentation errors. This is most clearly demonstrated on Atera, where despite the platform already providing whole-transcriptome coverage, cellpin further improved interpretability by suppressing residual technical noise, increasing transcriptional coherence within cell states, resolving biologically meaningful substructure, and improving downstream cell-type annotation. The value of cellpin extends well beyond recovering missing transcriptome coverage; it improves the biological coherence of spatial profiles even when full-transcriptome data is already available. Scalability is equally critical in this context as Atera can generate atlas-scale datasets in a single experiment. Existing methods would struggle with level of throughput, making cellpin’s lightweight design a practical necessity.

Several limitations of cellpin merit consideration. Benchmarking has focused primarily on 10x Genomics spatial platforms and MERSCOPE, while performance on technologies such as CosMx^41^ or seqFISH^42^ remains to be systematically evaluated, given differences in panel design, segmentation, and noise characteristics. Imputation quality is inherently bounded by the quality and completeness of the single-cell reference. Poorly annotated, low-coverage, biologically mismatched, or contamination-affected references, including those influenced by ambient RNA, dissociation artifacts or other technical noise, will limit transfer fidelity regardless of model architecture. Finally, while cellpin is highly scalable, training on very large references still requires suitable computational infrastructure, which may pose practical constraints for some users. As high-quality tissue atlases continue to expand in scope and resolution, the ceiling for cellpin performance should rise accordingly. Future extensions incorporating spatial priors or multi-omics integration are natural next steps for cellpin. Its ability to denoise and interpret large spatial datasets from a clean single-cell prior, without ever training on spatial data, positions cellpin as a robust and adaptable tool as spatial transcriptomic technologies continue to mature.

## Methods

### Cellpin overview

Cellpin imputes unmeasured genes in spatial transcriptomics by learning, from a scRNA-seq reference alone, a latent representation that can be inferred from either a full expression profile or a restricted gene panel. Each reference cell is represented as two aligned views: a full-gene vector over the retained reference genes and a panel vector containing only genes measured in the spatial assay. Training is performed exclusively on scRNA-seq cells. After training, spatial inference uses the panel encoder, a panel-based library-size encoder, and a shared decoder to reconstruct the retained reference gene space.

### Input representation and alignment

Cellpin operates directly on raw counts. Preprocessing identifies the intersection between reference and spatial genes, removes spatial genes absent from the reference, and reorders the overlapping panel genes to match their positions in the scRNA-seq reference. This ordering is enforced so that panel genes are presented consistently during training and inference. In our benchmark experiments, the retained reference space comprised the panel genes, along with 1,000 additional highly variable genes. However, we observed that cellpin performance remains stable with more than 15,000 reference genes (**Supplementary Table 2**).

### Model architecture

Cellpin uses two view-specific variational encoders and a shared decoder. The full encoder receives the retained reference profile and defines the reference latent geometry, whereas the panel encoder receives only the spatially measured genes and is trained to map into the same latent space. By default, both latent encoders use a hidden width of 1,024, 16 residual MLP blocks, and a 192-dimensional latent space. Each encoder consists of an input projection followed by residual feed-forward blocks with layer normalization, GEGLU activations, LayerScale, dropout, and linearly increasing DropPath regularization. Posterior means and variances are predicted from the final hidden state, with variances constrained by a softplus transform.

A separate library-size encoder predicts log-library size from panel genes alone. This branch uses the same hidden width but is intentionally shallow, with only a single residual block, and does not receive the Gaussian input perturbation used for the latent encoders.

The shared decoder reconstructs the retained full-gene profile from the latent representation and inferred library size. Its output likelihood follows a standard scVI^43^-style parameterization for count data, with negative binomial reconstruction and gene-specific global dispersion by default. However, cellpin couples this probabilistic output layer to a substantially higher-capacity encoder-decoder backbone than typically used, allowing the decoder to model more complex full-transcriptome structure while remaining compatible with panel-only inference.

### Two-stage training

Training proceeds in two stages. In Stage 1, the full encoder, shared decoder, and library encoder are trained on scRNA-seq data using a variational reconstruction objective on the retained full-gene profile. The latent state is inferred from the full view, whereas library size is inferred from the panel view. This stage establishes the reference latent space and trains the decoder to reconstruct the retained scRNA-seq gene space.

In Stage 2, the panel encoder is trained to recover the same latent structure from restricted panel input while preserving full-gene reconstruction. For each minibatch, the full encoder is evaluated without gradient updates and serves as a teacher. The panel encoder is then optimized using reconstruction, KL regularization, and latent-space matching to the teacher representation. By default, the decoder remains trainable during Stage 2, although a frozen-pretrained variant is also supported.

### Stage-2 loss and latent-space matching

For a minibatch of reference cells, the Stage-2 objective is

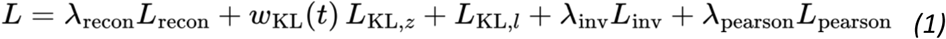

Here *L*_*recon*_ is the negative binomial reconstruction loss on the retained full-gene profile using the panel encoder and shared decoder; *L*_*KL,z*_ and *L*_*KL,l*_ regularize the cell-state and library-size latents; and *w*_*KL*_(*t*) use a linear KL warmup schedule. The invariance loss is described as:

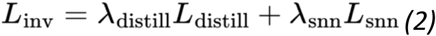

By default, *L*_*distill*_ is a mean-squared-error loss between the panel-derived and full-derived posterior means. Cellpin also supports a KL-based posterior distillation variant, but this is not the default configuration. The soft nearest-neighbor^44^ term *L*_*snn*_ is implemented as a symmetric cross-entropy loss over cosine-similarity logits between panel-derived and full-derived latent means, with same-cell pairs treated as positives. This encourages the panel encoder not only to match the teacher representation cell-by-cell but also to preserve the local neighborhood structure of the teacher-defined latent space within each minibatch.

### Training augmentations

To improve robustness to the domain gap between scRNA-seq reference data and spatial measurements, cellpin perturbs the input during training. First, panel counts are Poisson-resampled after multiplication by a randomly sampled efficiency factor to simulate reduced capture efficiency:

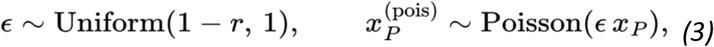

where *r* is the Poisson resampling rate ((*r*=0.1) by default). Second, contamination-style mixup is applied within each minibatch by blending each panel profile with that of a randomly selected cell:

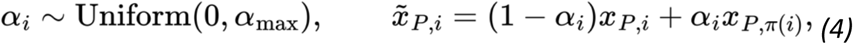

where *pi*(*i*) denotes a random permutation of cells in the minibatch and *α*_*max*_ = 0.1 by default. Third, Gaussian noise is added to the inputs of the two latent encoders during training:

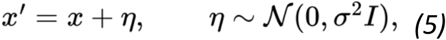

with *sigma* = 0.1 by default. This perturbation is applied to the full and panel latent encoders, but not to the library-size encoder, which is intended to estimate transcript abundance directly from observed panel counts.

### Library-size modeling

The library latent is modeled from panel genes alone in both training stages. Its prior is a univariate normal distribution with mean and variance estimated from log-transformed total counts in the training dataset. This separates the global count scale from the cellular state and ensures that the same library branch can be used at spatial inference time without access to full-gene profiles.

### Batch conditioning

Cellpin optionally conditions the decoder on categorical batch labels from the scRNA-seq reference. During spatial inference, when no matching batch label is available, the decoder receives a uniform soft one-hot vector across reference batches. Batch conditioning was disabled in benchmark experiments to ensure consistent comparison across methods, but can improve performance when integrating large or heterogeneous reference atlases.

### Spatial inference and imputation

After training, a spatial cell is processed by passing its measured panel counts through the panel encoder and the library-size encoder, followed by decoding into the retained reference gene space. Cellpin includes a safeguard that automatically detects and corrects panel-gene order mismatches between the training and inference datasets. By default, inference uses the posterior mean of the panel encoder for the embedding and the posterior mean of the library encoder for library size.

To improve robustness to noisy or partially corrupted panel measurements, cellpin performs Monte Carlo imputation by default. During inference, the measured panel is passed through the panel encoder *K* = 50 times. In each pass, a random subset of panel genes *p*_*drop*_ = 0.2 is temporarily masked prior to encoding, after which a latent sample is drawn from the panel posterior and decoded. The final imputed expression profile is obtained by averaging the predicted gene-wise means across stochastic forward passes:

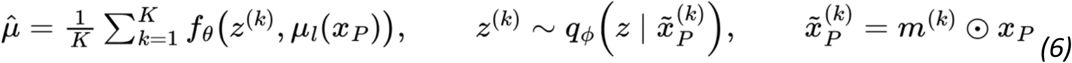

with

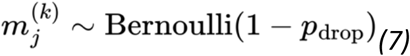

Cellpin additionally provides an internal normalization procedure for downstream log-transformed analyses. Rather than applying normalization and log-transformation directly to the averaged negative binomial mean, the method estimates the expectation of normalized, log-transformed counts by repeated sampling from the fitted negative binomial distribution:

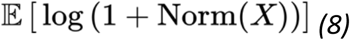

Where *Norm*(.) denotes either total-count normalization or area-based normalization when cell area is available. This corrects the bias that would arise from applying nonlinear normalization and log-transformation only after averaging. Optionally, imputed counts can also be returned as rounded integers (‘return_int = True’), which can provide a practically useful denoised count representation for downstream analyses.

### Label transfer

Cellpin includes a label-transfer function that annotates spatial observations in the learned latent space. Cellpin first represents both reference and spatial cells by their cellpin embeddings and then trains a distance-weighted k-nearest-neighbor classifier on the reference embedding using the known scRNA-seq cell type labels. Classifier performance can be assessed by stratified hold-out evaluation on the reference cells.

The trained classifier is then applied to the spatial embedding to assign cell type labels. For each spatial observation, cellpin returns both the predicted annotation and the corresponding maximum class probability as a confidence score. Optionally, observations with confidence below a user-defined threshold can be assigned to an “Unknown” category.

### Benchmark evaluation

#### Datasets and Preprocessing

We benchmarked gene expression prediction across a wide range of tissue contexts using both paired and unpaired single-cell RNA-sequencing (scRNA-seq) reference datasets together with spatial transcriptomics data (**Supplementary Table 1**). Unpaired reference datasets included out-of-batch atlas references to evaluate model robustness under realistic cross-dataset conditions. Spatial transcriptomics profiles were primarily generated using 10x Genomics Xenium technology, with an additional independently generated MERFISH dataset included as previously described^20^. For each dataset, analyses were restricted to the shared spatial gene panel together with the next 1,000 highly variable genes identified from the scRNA-seq reference dataset. scRNA-seq reference datasets containing more than 100,000 cells were randomly downsampled to 100,000 cells prior to model training, and Xenium slides were subsampled into square regions containing approximately 30,000–50,000 cells to avoid out-of-memory limitations in baseline methods. The breast cancer dataset^24^ and the 10x colorectal cancer dataset (**Supplementary Table 1**) were preprocessed following single-cell best-practice guidelines^45^, whereas the remaining datasets were used as provided as they had already undergone preprocessing. Raw count matrices were used both as model inputs and for evaluation.

#### Gene holdout benchmark

To evaluate spatial gene expression prediction in a controlled and unbiased setting, we implemented a five-fold gene holdout benchmark. Prior to model training, we defined a set of eligible holdout genes consisting of genes shared between the scRNA-seq and spatial datasets that exhibited non-zero mean expression and non-zero variance in both modalities. This filtering step ensured compatibility across all benchmarked methods, including approaches that internally exclude constant-expression genes.

For each fold, 50 genes were randomly sampled without replacement from the eligible gene pool using a fixed random seed (42), resulting in 250 unique holdout genes per dataset across all folds. Holdout genes were removed from the spatial input during training, and each method was tasked with predicting their expression across all retained spatial locations. Predicted expression values were subsequently compared against the measured spatial transcriptomic profiles, which served as ground truth.

#### Baseline methods

Cellpin was benchmarked against five previously published methods for spatial gene expression prediction as well as a foundational model evaluated in both zero-shot and fine-tuned settings: SpaGE, gimVI, Tangram, scENVI, stDiff, scConcept, and scConcept+. All methods were trained and evaluated on identical benchmark splits using published default parameters. Implementations for SpaGE, gimVI, and Tangram were obtained through the SpatialBenchmarking framework^12^, while the remaining methods were run following their original published implementations and default parameters. Cellpin was evaluated under the same conditions using the default model settings (latent dimension: 192, batch size: 256, encoder dimensions: 16) without providing batch labels or additional covariates. Cellpin was pretrained for 50 epochs and subsequently trained for up to 80 epochs with early stopping enabled. Tangram was executed in CPU mode, as GPU execution consistently resulted in out-of-memory errors on NVIDIA A40 GPUs.

#### Evaluation metrics

Prediction performance was evaluated exclusively on held-out genes and computed independently for each gene across all spatial cells using four complementary metrics capturing distinct aspects of predictive accuracy. Pearson correlation coefficient (r) was used to quantify the linear relationship between predicted and observed expression values, whereas Lin’s concordance correlation coefficient (CCC) additionally penalized systematic deviations from the identity line and therefore captured both correlation and absolute agreement. Root mean square error (RMSE) was computed on z-score-normalized expression values to account for differences in gene-specific expression magnitude and dynamic range, enabling comparisons across genes with highly variable abundance levels. To further assess how well models recovered the overall expression landscape, Jensen–Shannon (JS) divergence was calculated between predicted and observed expression distributions after normalization to probability distributions, providing a symmetric measure of distributional similarity. In addition, we computed a per-cell cosine similarity across all combined held-out genes for each cell to quantify agreement between predicted and observed transcriptomic profiles at the cellular level (250 genes per cell). Per-cell cosine similarities were then aggregated across datasets for global comparisons. Finally, to assess whether models preserved gene-specific variability across the sample, we compared the variance and mean of predicted and ground-truth expression for each held-out gene and quantified their agreement using Pearson correlation.

#### General analysis

Analysis and processing of spatial samples was performed using scanpy ^46^, anndata^39^, squidpy^47^, and spatialdata^40^ and following single cell best practices^45^.

#### Sample-level representation analysis

To assess sample-level trajectory, each donor sample was split into three pseudoreplicates by randomly assessing cells to a pseudoreplicate. Per-sample quality metrics, cell type composition, and pseudobulk representations were computed using patpy (v 0.9.2)^32^. Pseudobulk embeddings were constructed from the cellpin latent space by aggregating cell embeddings across cells per pseudoreplicate using patpy.tl.Pseudobulk. Pairwise sample distances were computed in this aggregated embedding space and visualized via UMAP. Disease progression trajectories were inferred using diffusion pseudotime (DPT) rooted at the least-affected sample.

#### ResolVI imputation

Imputation with ResolVI was performed using the published implementation and default model settings, following the authors’ recommended workflow. The model was trained for 100 epochs, as recommended in the original documentation and tutorial.

#### Noisy genes fraction

To assess noise, we estimated gene detection probabilities for each cell type as the fraction of cells in which each gene was detected, using Laplace-smoothed estimates of the probability of a gene to be detected in a certain cell type. A gene was considered “noisy” in a given cell if it was detected despite having a low probability of detection within that cell’s annotated cell type. For each cell, we computed the noisy fraction, defined as the proportion of detected genes classified as noisy. Lower noisy fractions indicate improved cell type specificity and reduced stochastic noisy expression.

## Code and data availability

The source code for cellpin is publicly available at [GitHub](https://github.com/Saur-Lab/cellpin), with documentation hosted on [Read the Docs](https://cellpin.readthedocs.io/). All datasets used in this study are publicly available and detailed in **Supplementary Table 1**.

## Acknowledgments

The study was supported by the German Cancer Consortium (DKTK) and DKTK Joint Funding, the DKFZ-MOST cooperation program (Ca-217), the Deutsche Forschungsgemeinschaft (SFB 1371 Project-ID 395357507 P12 to D.S.; DFG SA 1374/8-1 Project-ID 515991405 to D.S.; DFG SA 1374/7-1 Project-ID 515571394 to D.S., F.T.; DFG SA 1374/6-1 Project-ID 458890590 to D.S.); Deutsche Krebshilfe (#70117118 DEFEAT-PDAC of the German Pancreatic Cancer Alliance to D.S., F.T.; #70115743 to D.S.; #70116843 to D.S.).

## Author contributions

P.P. and D.L. contributed equally to this work. P.P. and D.L. conceptualized the study, developed the Cellpin software framework, and performed the data analyses. P.P., D.L., and D.W. generated the figures and wrote the original draft of the manuscript. M.B. implemented and ran the scConcept baseline for the benchmarking evaluation. D.S. acquired funding. The study was jointly supervised by M.L., F.J.T. and D.S. M.L., F.J.T. and D.S. proofread, reviewed, and edited the manuscript. All authors proofread and approved the final form of the manuscript.

