## Supplementary Figures for "Cellpin enables reference-based imputation and denoising of spatial transcriptomes"

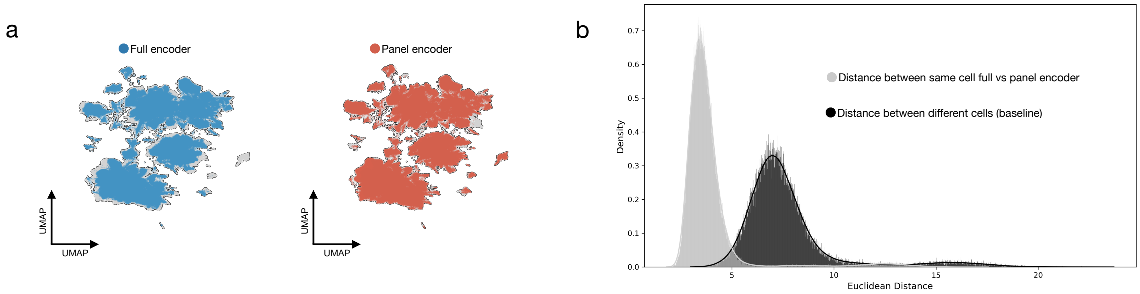


**Supplementary Figure 1. a)** Core GBMap scRNA-seq atlas[24] (**Supplementary Table 1**) embedded using the cellpin full encoder with 1,450 reference genes (left) and the panel encoder with training augmentations using 450 genes (right), showing strong overlap between different embeddings of the same cells. **b)** Distribution of Euclidean distances between the same cells encoded with the cellpin full encoder and panel encoder with augmentation, compared with distances between different cells as a baseline.

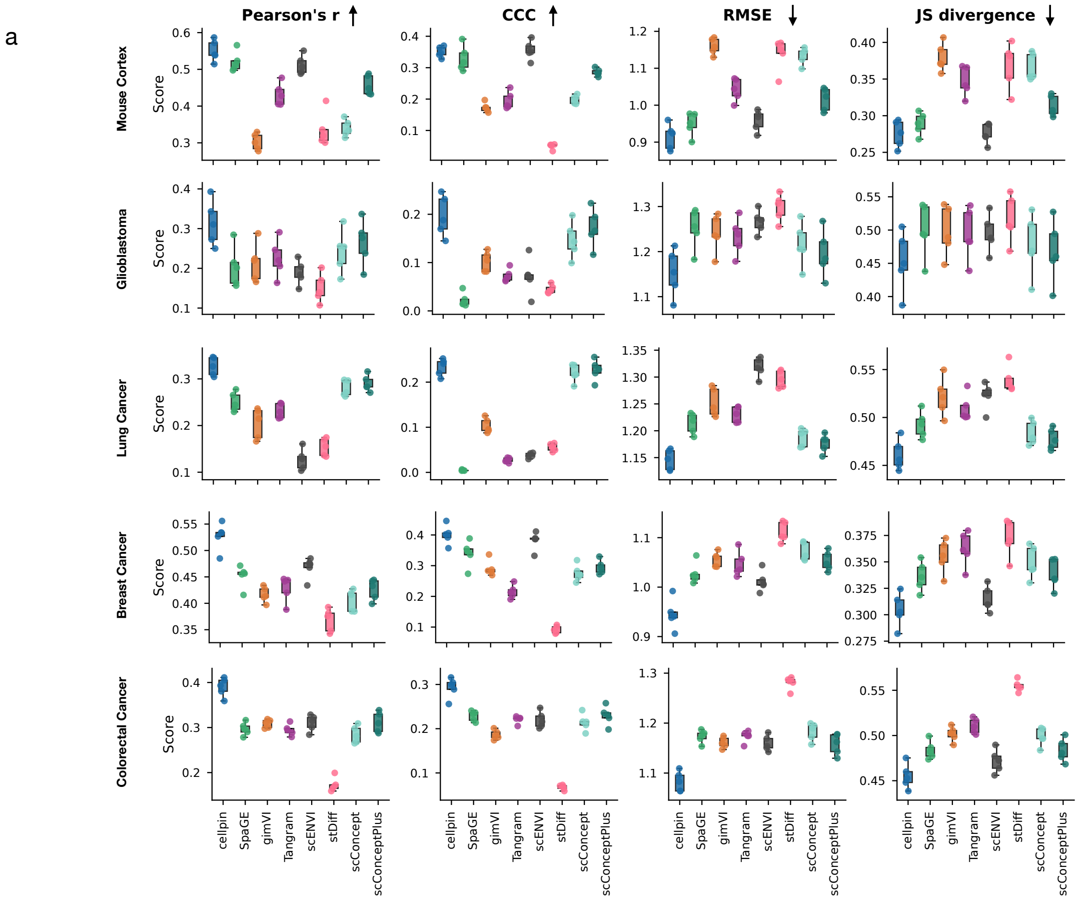


**Supplementary Figure 2. a)** Mean held-out Pearson correlation (Pearson's *r*), Lin's concordance correlation coefficient (CCC), root mean square error (RMSE), and Jensen–Shannon divergence (JS) across five datasets (**Supplementary Table 1**). Each dot represents one of five cross-validation folds, each containing 50 unique held-out genes (250 genes per dataset total).

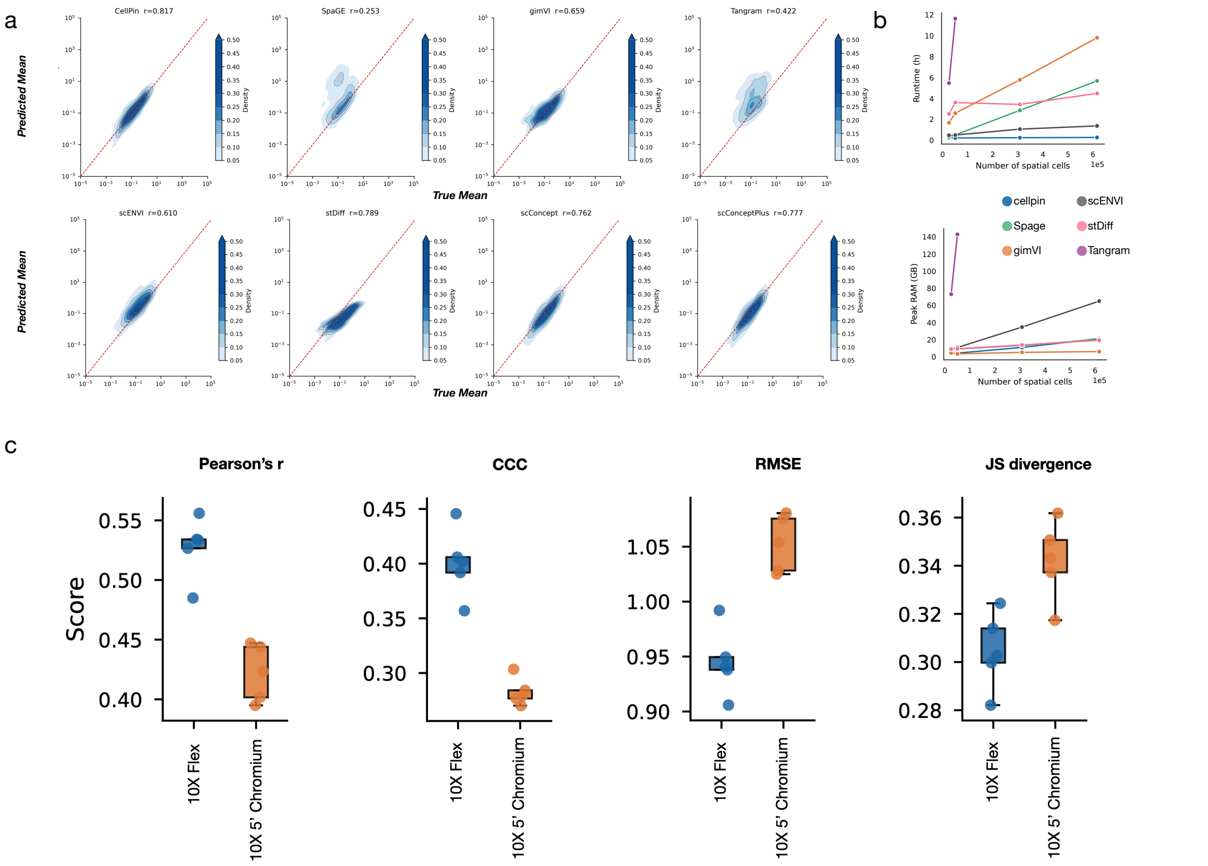


**Supplementary Figure 3. a)** True versus predicted mean expression for all held-out genes across 5 datasets, with Pearson correlation (r) between true and predicted mean annotated for each method. **b)** Runtime (in hours (h)) (top) and peak RAM usage (in giga-byte (GB)) for training cellpin, SpaGE, gimVI, scENVI, stDiff, and Tangram with 100,000 single-cell RNA reference cells and 25,000, 50,000, 307,762 (full slide) and 615524 (duplicated full slide) spatial cells from the 10X Colorectal Cancer dataset (**Supplementary Table 1**). Tangram consumed >250 GB of RAM using the full slide or more and was therefore aborted. scConcept (foundation model) not shown. **c)** 5-fold evaluation of the Matched Breast Cancer data using either 10X Flex or 10X 5' Chromium as reference (**Supplementary Table 1**). Pearson correlation (Pearson's r), Lin's concordance correlation coefficient (CCC), root mean square error (RMSE) and Jensen-Shannon divergence (JS) shown. Each fold consists of 50 non-overlapping genes.

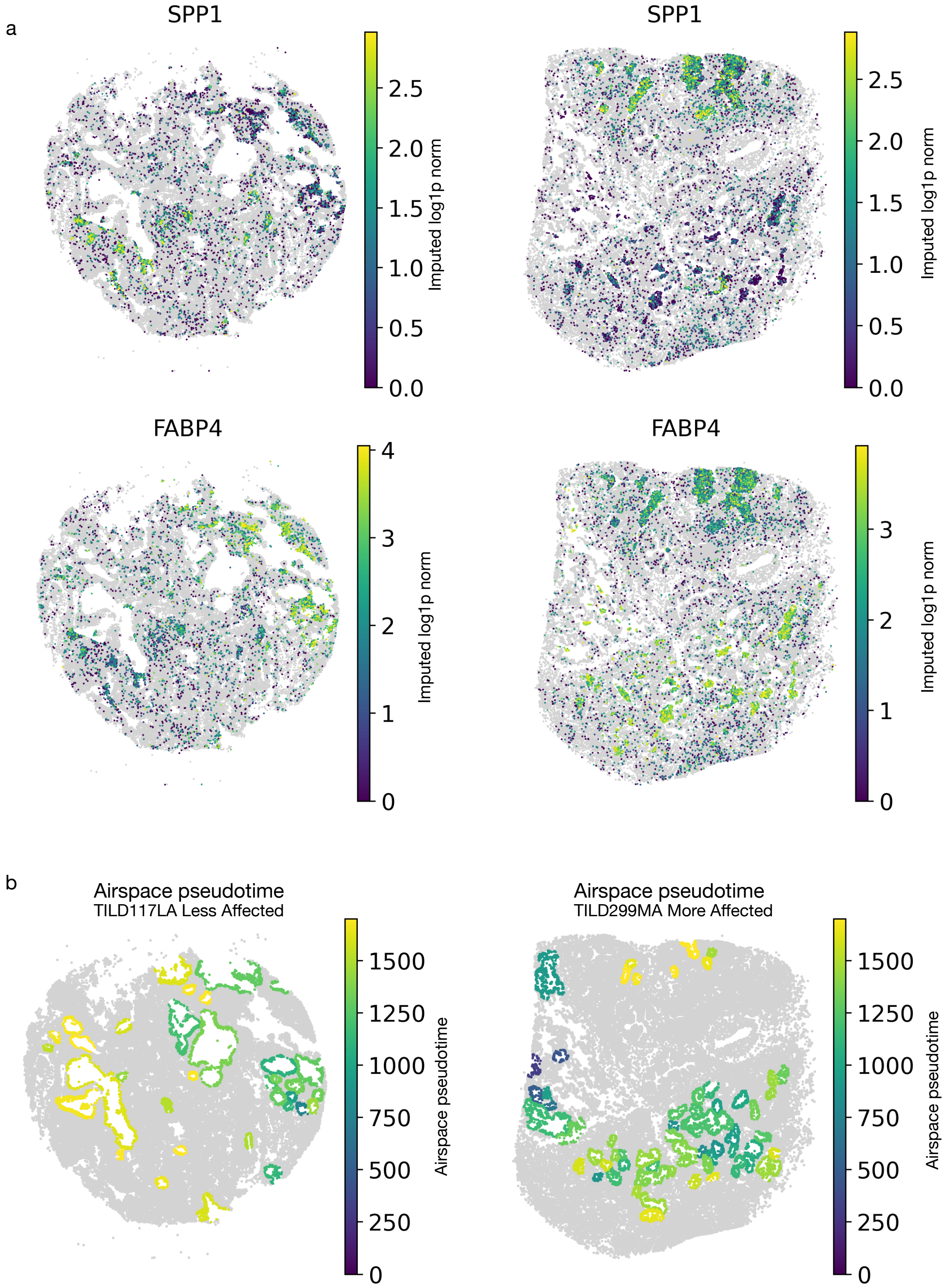


**Supplementary Figure 4. a)** Spatial expression of cellpin-imputed *SPP1* and *FABP4* in macrophages in two representative lung tissue sections**. b)** Spatial map of airspace pseudotime (*Lumen_rank*) in the corresponding sections, as introduced in the original manuscript[27].

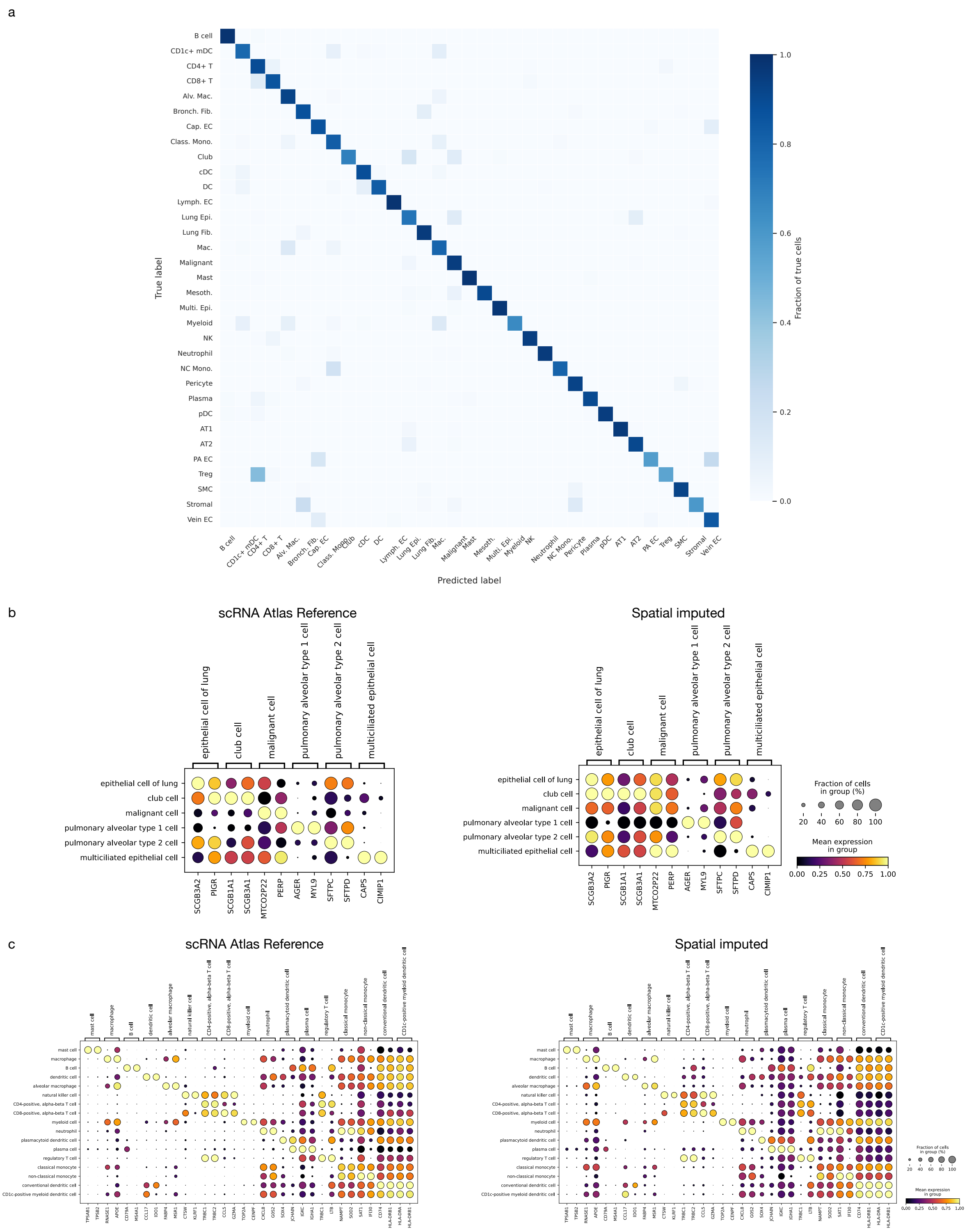


**Supplementary Figure 5. a)** Performance of the kNN classifier on the held-out test set of the single cell reference[25]. **b)** Dot plots showing mean expression of epithelial cell type marker genes in the scRNA-seq reference atlas (left) and cellpin-imputed Xenium data (right). Marker genes were identified by differential expression analysis in the reference atlas. **c)** Dot plots showing mean expression of immune cell type marker genes in the scRNA-seq reference atlas (left) and cellpin-imputed Xenium data (right). Marker genes were identified by differential expression analysis in the reference atlas.

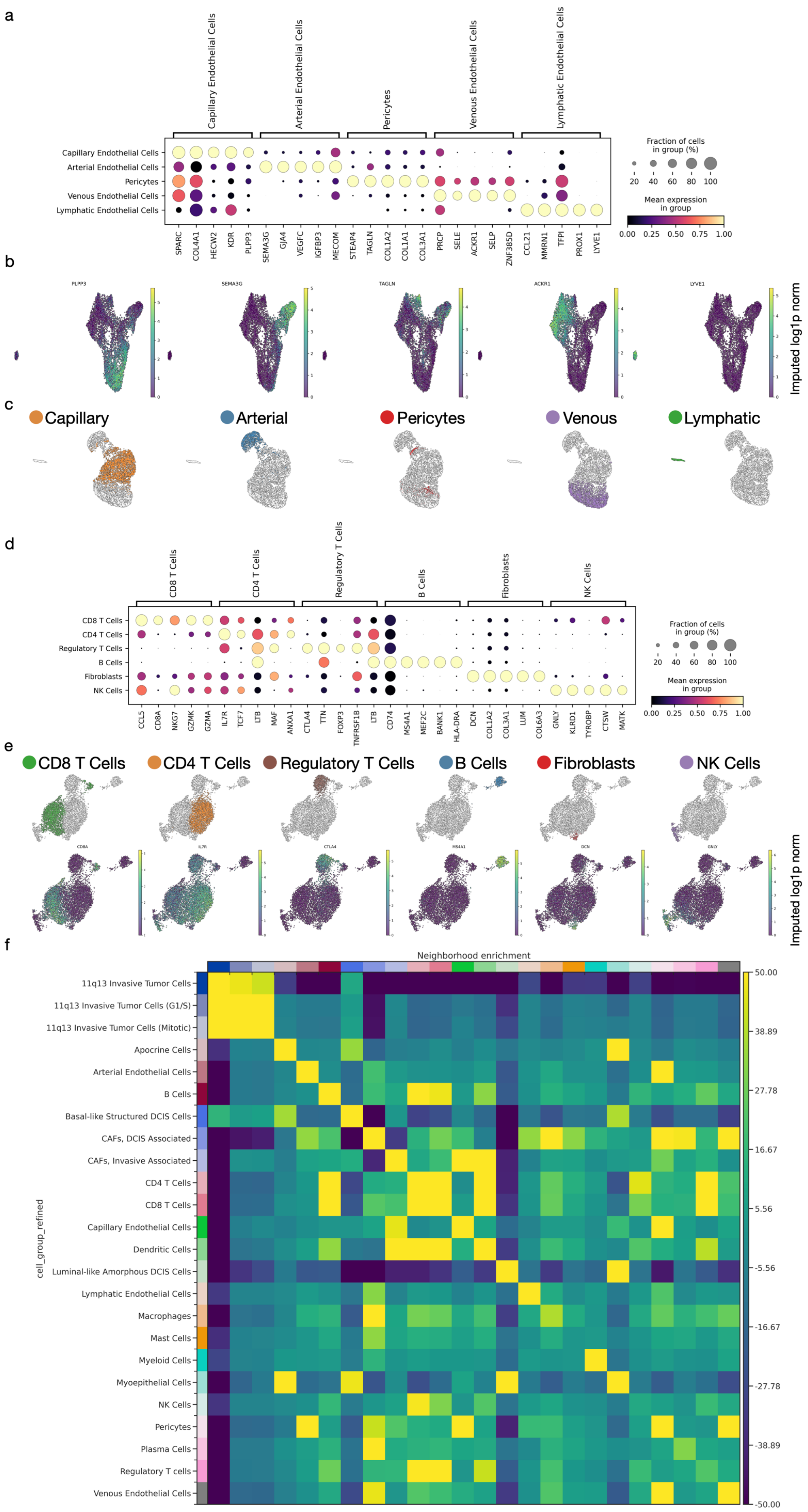
a
Capillary Endothelial Cells
Arterial Endothelial Cells
Pericytes
Venous Endothelial Cells
Lymphatic Endothelial Cells
SPARC
COL4A1
HECW2
KDR
PLPP3
SEMA3G
GJA4
VEGFC
IGFBP3
MECOM
STEAP4
TAGLN
COL1A2
COL1A1
COL3A1
PRCP
SELE
ACKR1
SELP
ZNF385D
CCL21
MMRN1
TFPI
PROX1
LYVE1
Fraction of cells in group (%)
20
40
60
80
100
Mean expression in group
0.00
0.25
0.50
0.75
1.00
b
PLPP3
SEMA3G
TAGLN
ACKR1
LYVE1
Imputed log1p norm
c
Capillary
Arterial
Pericytes
Venous
Lymphatic
d
CD8 T Cells
CD4 T Cells
Regulatory T Cells
B Cells
Fibroblasts
NK Cells
CCL5
CD8A
NKG7
GZMK
GZMA
IL7R
TCF7
LTB
MAF
ANXA1
CTLA4
TTN
FOXP3
TNFRSF1B
LTB
CD74
MS4A1
MEF2C
BANK1
HLA-DRA
DCN
COL1A2
COL3A1
LUM
COL6A3
GNLY
KLRD1
TYROBP
CTSW
MATK
Fraction of cells in group (%)
20
40
60
80
100
Mean expression in group
0.00
0.25
0.50
0.75
1.00
e
CD8 T Cells
CD4 T Cells
Regulatory T Cells
B Cells
Fibroblasts
NK Cells
CD8A
IL7R
CTLA4
MS4A1
DCN
GNLY
Imputed log1p norm
f
Neighborhood enrichment
cell_group_refined
11q13 Invasive Tumor Cells
11q13 Invasive Tumor Cells (G1/S)
11q13 Invasive Tumor Cells (Mitotic)
Apocrine Cells
Arterial Endothelial Cells
B Cells
Basal-like Structured DCIS Cells
CAFs, DCIS Associated
CAFs, Invasive Associated
CD4 T Cells
CD8 T Cells
Capillary Endothelial Cells
Dendritic Cells
Luminal-like Amorphous DCIS Cells
Lymphatic Endothelial Cells
Macrophages
Mast Cells
Myeloid Cells
Myoepithelial Cells
NK Cells
Pericytes
Plasma Cells
Regulatory T cells
Venous Endothelial Cells
50.00
38.89
27.78
16.67
5.56
-5.56
-16.67
-27.78
-38.89
-50.00

**Supplementary Figure 6. a)** Dot plot showing mean expression of marker genes in re-annotated blood vessel cell populations in Cellpin-imputed Atera data. **b)** Representative uniform manifold approximation and projection (UMAP) visualizations showing expression of blood vessel cell type marker genes in the imputed latent space. **c)** UMAP derived from the PCA computed over the imputed gene space shows a separation of the main cell types individuated from the cellpin embedding. **d)** Dot plot showing mean expression of marker genes in the re-annotated T lymphocyte clusters in Cellpin-imputed Atera data. **e)** Representative UMAP visualizations of the re-annotated T lymphocyte cell types (top) and corresponding marker gene expression in the imputed latent space (bottom). **f)** Spatial neighborhood enrichment analysis showing colocalization of capillary endothelial cells and invasion-associated CAFs.
