## Supplementary Figures for "Cellpin enables reference-based imputation and denoising of spatial transcriptomes"

**Supplementary Table 1: Overview of all datasets used in the study**

| Dataset | Technology | Origin | Dataset Link | Manuscript | Use-case | Figure |
|---|---|---|---|---|---|---|
| Mouse_Cortex | MERFISH | ENVI Tutorial | https://dp-lab-data-public.s3.amazonaws.com/ENVI/st_data.h5ad | https://doi.org/10.1038/s41586-021-03705-x | Benchmark | Figure 2 |
| Mouse_Cortex | scRNA | ENVI Tutorial | https://dp-lab-data-public.s3.amazonaws.com/ENVI/sc_data.h5ad | https://doi.org/10.1038/s41586-021-03500-8 | Benchmark | Figure 2 |
| Janesick_Breast Cancer_Sample_1_Replicate_1 | Xenium | 10X Genomics Datasets | https://www.10xgenomics.com/products/xenium-in-situ/preview-dataset-human-breast | https://doi.org/10.1038/s41467-023-43458-x | Benchmark, Graphical Abstract | Figure 1, Figure 2 |
| Janesick_Breast Cancer_FRP | scRNA | 10X Genomics Datasets | https://www.10xgenomics.com/products/xenium-in-situ/preview-dataset-human-breast | https://doi.org/10.1038/s41467-023-43458-x | Benchmark, Graphical Abstract | Figure 1, Figure 2 |
| 10X_Colorectal Cancer_P1 | Xenium | 10X Genomics Datasets | https://www.10xgenomics.com/platforms/visium/product-family/dataset-human-crc | https://doi.org/10.1038/s41588-025-02193-3 | Benchmark | Figure 2 |
| 10X_Colorectal Cancer_FRP_aggregated | scRNA | 10X Genomics Datasets | https://www.10xgenomics.com/platforms/visium/product-family/dataset-human-crc | https://doi.org/10.1038/s41588-025-02193-3 | Benchmark | Figure 2 |
| Janesick_Breast Cancer_5prime | scRNA | 10X Genomics Datasets | https://www.10xgenomics.com/products/xenium-in-situ/preview-dataset-human-breast | https://doi.org/10.1038/s41467-023-43458-x | Reference Quality Assesement | Supplementary Figure 3 |
| 10X_Glioblastoma | Xenium | 10X Genomics Datasets | https://www.10xgenomics.com/datasets/ffpe-human-brain-cancer-data-with-human-immuno-oncology-profiling-panel-and-custom-add-on-1-standard | n/a | Benchmark | Figure 2 |

| | | | | | | |
|---|---|---|---|---|---|---|
| Core_GBMap | scRNA | Cell x Gene | https://cellxgene.cziscience.com/collections/999f2a15-3d7e-440b-96ae-2c806799c08c | https://doi.org/10.1093/neuonc/noaf113 | Benchmark | Figure 2 |
| 10X_Lung_Cancer | Xenium | 10X Genomics Datasets | https://www.10xgenomics.com/datasets/preview-data-ffpe-human-lung-cancer-with-xenium-multimodal-cell-segmentation-1-standard | n/a | Benchmark | Figure 2, Figure 4 |
| Salcher_Atlas | scRNA | Cell x Gene | https://cellxgene.cziscience.com/collections/edb893ee-4066-4128-9aec-5eb2b03f8287 | https://www.cell.com/cancer-cell/fulltext/S1535-6108(22)00499-8 | Benchmark, Figure Reference | Figure 2, Figure 4 |
| LungFibrosis_Xenium | Xenium | GEO | https://www.ncbi.nlm.nih.gov/geo/query/acc.cgi?acc=GSE250346 | https://doi.org/10.1038/s41588-025-02080-x | Reproduce Findings, Analysis | Figure 3 |
| HCLA | scRNA | Cell x Gene | https://cellxgene.cziscience.com/collections/6f6d381a-7701-4781-935c-db10d30de293 | https://doi.org/10.1038/s41591-023-02327-2 | Reference Figure 3 | Figure 3 |
| ChenBreastCancer Atlas | scRNA | Cell x Gene | https://cellxgene.cziscience.com/collections/9432ae97-4803-4b9f-8f64-2b41e42ad3cb | https://doi.org/10.1093/nargab/lqaf217 | Reference Atera | Figure 5 |
| Breast_Cancer_Atera | Atera | 10X Genomics Datasets | https://www.10xgenomics.com/datasets/atera-wta-ffpe-human-breast-cancer | n/a | Use-case Atera | Figure 5 |

Overview of all datasets used in this study

### Supplementary Table 2: Runtime and memory usage of imputation methods

| scRNA + spatial | cellpin [Time in s] | scENVI [Time in s] | gimVI [Time in s] | Tangram (CPU) [Time in s] | stDiff [Time in s] | spaGE [Time in s] |
|---|---|---|---|---|---|---|
| 100k + 25k | **790,10** | 1752,20 | 6082,60 | 19715,60 | 9128,10 | 1083,10 |
| 100k + 50k | **826,10** | 1845,20 | 9362,20 | 41887,10 | 13004,50 | 1914,20 |
| 100k + 307kk | **922,10** | 3879,50 | 20838,00 | aborted >250GB RAM | 12377,60 | 10333,50 |
| 100k + 615k | **1023,10** | 4982,6 | 35289,50 | aborted >250GB RAM | 16171,00 | 20467,80 |

| scRNA + spatial | cellpin [RAM in GB] | scENVI [RAM in GB] | gimVI [RAM in GB] | Tangram (CPU) [RAM in GB] | stDiff [RAM in GB] | spaGE [RAM in GB] |
|---|---|---|---|---|---|---|
| 100k + 25k | **3,777** | 8,795 | 4,479 | 73,087 | 9,014 | 9,457 |
| 100k + 50k | **4,246** | 10,791 | 4,342 | 143,007 | 9,360 | 9,065 |
| 100k + 307kk | 10,904 | 34,681 | **5,155** | aborted >250GB RAM | 13,732 | 13,265 |
| 100k + 615k | 21,168 | 64,999 | **6,051** | aborted >250GB RAM | 19,269 | 20,304 |

Runtime seconds and peak RAM usage (in giga-byte) for training cellpin, SpaGE, gimVI, scENVI, stDiff, and Tangram with 100,000 single-cell RNA reference cells and 25,000, 50,000, 307,762 (full slide) and 615524 (duplicated full slide) spatial cells from the 10X Colorectal Cancer dataset. Tangram consumed >250 GB of RAM using the full slide or more and was therefore aborted. scConcept (foundation model) not shown.

### Supplementary Table 3: Heldout pearson correlation with increasing number of reference genes

| n_genes | Mean Pearson | Median Pearson | std |
|---|---|---|---|
| 407 | **0.5336866179** | 0.550244985 | **0.2089985494** |
| 907 | 0.5242936124 | 0.5422738658 | 0.2096492359 |
| 1307 | 0.5291377093 | 0.5462255727 | 0.2123485611 |
| 2307 | 0.5311821782 | 0.5486410036 | 0.2101290463 |
| 5307 | 0.5210870542 | 0.547468937 | 0.2108383885 |
| 10307 | 0.5271300317 | **0.552693774** | 0.2108669016 |
| 15307 | 0.5183415765 | 0.5460317408 | 0.2126499276 |

Imputation performance was evaluated across six gene reference sizes (407–15,307 genes) using the Janesick et al. dataset. Performance is reported as the mean and median Pearson correlation between predicted and observed expression values across 50 held-out genes. Standard deviation (s.d.) was calculated across the held-out genes. n_genes denotes the total number of genes available to the model as reference input during training.
